# Identity-stable multi-animal tracking using bidirectional segmentation with object-level memory

**DOI:** 10.1101/2025.09.22.677949

**Authors:** Shuyu Wang, Kara Quine, Audrey Jordan, Shreya Dasari, Devanand S. Manoli, Christoph Kirst

## Abstract

Tracking animal behavior in naturalistic settings is essential for understanding social dynamics and their neural underpinnings. Pose estimation methods can produce accurate keypoints using framewise inference. However, post hoc tracking steps often struggle to maintain consistent identity over time, particularly during close and rapid social interactions between visually similar animals. We present a pipeline for bidirectional video object segmentation (VOS) to correct identity swaps with much less manual annotation effort, addressing the prohibitive cost of identity correction of pose estimation data in long recordings and large cohorts. Our approach makes use of a state-of-the-art VOS algorithm, Cutie, which leverages both pixel- and object-level representations across multiple memory timescales. By comparing segmentation masks from independent forward and reverse inference runs, we identify localized zones of disagreement and flag them for manual review. When applied to more than 160 hours of dyadic vole interaction videos, our method reduces identity swaps by two orders of magnitude compared to typical pose estimation workflows and requires review of less than 0.3% of frames per video to achieve identity error-free segmentation masks and aligned keypoints. Our approach generalizes to social interactions involving three or more animals, with scalability constrained primarily by behavioral complexity (e.g., complete occlusion of multiple individuals). Our method enables scalable, long-term tracking of unmarked animals in group settings and provides a practical foundation for more naturalistic studies of social behavior. To lower the barrier for researchers facing similar tracking challenges, we provide an accessible graphical user interface for general use.

## INTRODUCTION

Quantitative measurement of naturalistic behavior is essential for neuroscience,^1–3^ ethology,^1–4^ and ecology.^5^ Robust behavioral quantification enables researchers to capture natural variation as well as changes induced by experimental perturbations. In non-human species that cannot communicate verbally with experimenters, behavior arguably represents one of the most critical observable phenotypes. In recent years, markerless pose estimation algorithms such as SLEAP^6^, multi-animal DeepLabCut (maDLC)^7^, and DANNCE^8–10^ have transformed the study of animal behavior by enabling high-resolution tracking of movement with impressive accuracy and generalizability. A wide range of tools now leverage tracking data to extract behavioral structure through both supervised classification of known behaviors and unsupervised discovery of novel motifs.^1,11–21^

However, investigating social behavior – particularly over extended timescales – poses distinct challenges. Social behavior is highly dynamic, shaped by internal state,^22^ experience,^23^ and environmental context.^24,25^ It often unfolds over minutes to hours or longer. Clinically relevant phenomena such as isolation-induced distress^26,27^ and disruptions in social attachment^28^ depend on behavior that evolves gradually, making long-term, continuous tracking essential. Yet, achieving reliable identity tracking across prolonged interactions and variable conditions remains difficult, especially in species with limited visual identifiers or complex patterns of social behavior.

Prairie voles are a socially monogamous rodent species and a leading model for studying the neurobiology of social attachment and complex social behaviors. Unlike mice, which display social novelty preference,^29^ prairie voles form pair bonds characterized by strong partner preference,^30–32^ distress upon separation,^26,27^ and selective rejection of unfamiliar opposite-sex conspecifics.^30–32^ These behaviors are often sustained over the animals’ lifetime, making them especially well-suited for studying long-term social relationships. From a computational perspective, however, their uniform fur, highly deformable bodies, and short, often occluded tails complicate pose estimation and increase the likelihood of identity swaps. Their behavioral repertoire also includes prolonged interactions at close range with both affiliative and agonistic characteristics. Together, these attributes demand identity-stable tracking over extended periods, as even a single swap can obscure meaningful, often sex-specific, behavioral patterns.

Physical indicators such as body markings and RFIDs have historically been used to facilitate identity tracking,^33–35^ but they are limited by the need for animal handling, potential behavioral artifacts, and low tracking resolution.^36–40^ Existing identity-tracking algorithms perform well when individuals have stable, distinctive appearance features, but their accuracy declines for species with little visual differentiation and frequent physical contact.^41–46^ These limitations motivate the use of pose estimation frameworks such as SLEAP and maDLC, which localize keypoints, assemble them into individuals based on part affinity fields, and then apply post hoc identity assignment. In these frameworks (currently SLEAP v1.4.1 and maDLC v2.2), keypoints are detected and assembled independently on each frame, with identities subsequently linked across frames using appearance, proximity, and movement statistics.^6,7^ While this strategy supports efficient inference and avoids propagating detection errors, it does not exploit spatiotemporal continuity during inference. As a result, even these state-of-the-art tools are vulnerable to identity instability, particularly during prolonged physical interactions between visually similar animals.

To address these challenges, we developed a pipeline for bidirectional video object segmentation (VOS) powered by the Cutie algorithm^47^, which leverages spatiotemporal memory to achieve high-fidelity tracking of animal shape while substantially reducing identity swaps. VOS is inherently directional and history-dependent: forward inference from the start of a video and reverse inference from the end of a video yield two independently generated sets of masks.

We identify zones of disagreement (ZOD) by computing the intersection over union between forward and reverse segmentations, highlighting regions where the two inferences diverge. These ZODs reveal potential segmentation errors that can be resolved to generate consensus, identity-preserved masks. Beyond producing this complementary source of tracking information, our approach substantially lowers the burden of correcting identity swaps in pose estimation, reducing the required manual effort by multiple orders of magnitude. Finally, we will make available our code base, graphical user interface, and example materials.

## RESULTS

Our bidirectional Cutie-based video object segmentation pipeline combines (1) user-prompted keyframe mask generation, (2) segmentation inference, (3) targeted correction of mask inconsistencies, and (4) automated alignment of pose estimation keypoints to identity-validated masks (Fig. 1A). We first quantify the prevalence and difficulty of identity errors in pose-tracked data and then detail how each stage of our pipeline addresses these challenges. Altogether, this approach enables users to resolve identity errors at the level of segmentation masks – where they are rarer and easier to correct – rather than within pose estimation data, resulting in two synchronized, high-quality data streams: masks and keypoints.

**Figure 1:**
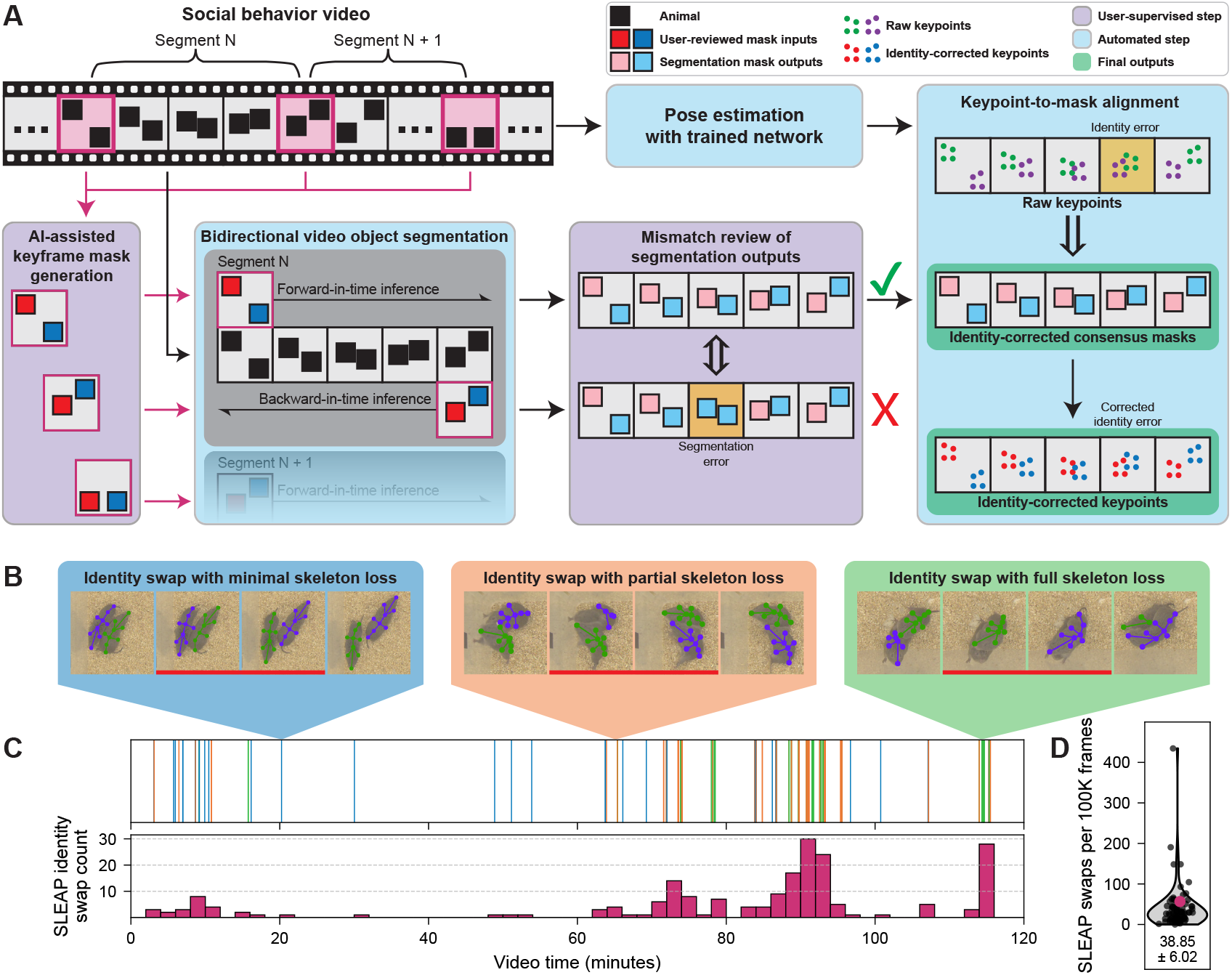
Bidirectional video object segmentation resolves identity swaps in dyadic pose estimation. A. Schematic of our pipeline for generating high-fidelity segmentation masks and correcting identity errors in pose estimation with minimal user input. The workflow includes: (1) user-prompted keyframe mask generation from Segment Anything, (2) bidirectional video object segmentation (VOS), (3) targeted resolution of segmentation mismatches via Zone of Disagreement (ZOD) review, and (4) automated alignment of pose estimation keypoints to identity-validated segmentation masks. Color-coded modules are explained in the legend (upper right). B. Identity swaps in pose estimation data fall into three categories based on the degree of keypoint loss: (1) minimal skeleton loss, where 8 or more keypoints are detected but assigned to the wrong individual (blue); (2) partial skeleton loss with 1-7 out of 11 keypoints detected (orange); and (3) full skeleton loss (emerald). Keypoint dropout often results from occlusion, atypical posture, or motion blur. For example, vole tails are short and frequently occluded, resulting in unreliable detection of the tail base and tail tip. Red bars indicate pairs of frames between which swaps have occurred. C. Distribution of manually annotated swaps in a representative two-hour video. Top: time series of swaps colored by type (see Fig. 1B). Bottom: histogram of swap counts in non-overlapping two-minute bins. See Supp. Fig. 1B for swap densities of five manually annotated videos. D. Distribution of SLEAP identity swap counts across 83 two-hour recordings (360,000 frames each) after automated alignment to Cutie-generated identity-preserved segmentation masks. The representative video from Fig. 1C is highlighted in magenta.

### Identity swaps in keypoint-based tracking hinders long-term social behavior analysis

To characterize social behavior in prairie voles, we first trained a SLEAP network on over 2,400 manually annotated frames captured from a top-view camera, with keypoints labeled for two interacting animals (see Methods). The network achieved pose estimation performance similar to published benchmarks for mice, including a comparable frequency of identity swaps after post hoc tracking (Fig. 1D).^6^ Identity errors fell into three categories (Fig. 1B): those involving fully visible skeletons (blue inset), partial skeleton loss (orange inset), or complete skeleton loss (emerald inset). These swaps occurred sporadically throughout the recordings (Fig. 1C, Supp. Fig. 1B). They occurred more frequently in frames with compromised pose estimation (Supp. Fig. 1C), which are typically associated with animal-on-animal occlusion, nonstandard postures, or extreme locomotor states (such as fighting). We tested a range of identity tracking algorithms and parameter configurations within SLEAP, but none consistently prevented identity swaps across long recordings (Supp. Fig. 1A, see Methods).

To correct identity swaps in SLEAP tracking data, we explored several post hoc strategies. Manual review of identity labels on a frame-by-frame basis proved prohibitively time-consuming, especially for hours-long recordings. As a heuristic, we computed inter-frame displacement for each track’s keypoints normalized by the keypoint count (Supp. Fig. 1C,Di).^7^ This approach flagged many swaps involving large inter-frame skeleton displacements, particularly those with substantial skeleton preservation of both animals (Fig. 1B blue inset).

However, it failed to flag swaps when inter-frame displacement was within the typical range (Supp. Fig. 1C). Identity swaps were also more frequent in frame pairs with fewer shared detected keypoints, indicating that incomplete detections compromise identity continuity (Supp. Fig. 1C,Div). Thus, ranking frame pairs by average displacement per keypoint from highest to lowest enriched for many swaps but did not yield an efficient search strategy overall: the long tail of swaps distributed across the ranking spectrum could not be localized without exhaustive inspection (Supp. Fig. 1Di). We also tested alternative metrics, including SLEAP’s built-in tracking scores, but these too failed to locate identity swap events reliably (Supp. Fig. 1Dii,Diii).^6^ Together, these findings highlighted the limitations of current post hoc tracking methods and motivated the development of an alternative inference strategy that preserves identity through fundamentally different principles.

### Bidirectional inference with Cutie, a video object segmentation network

We reasoned that machine learning models with built-in memory mechanisms would be less prone to identity confusion over time. Based on the hypothesis that shape information would enable detailed characterization of social touch, we sought a segmentation model capable of shape estimation to complement the pose estimation outputs of SLEAP inference.^48^ To this end, we selected Cutie, a video object segmentation (VOS) network that integrates bottom-up pixel-level and top-down object-level memory across multiple timescales (schematic of original architecture adapted in Supp. Fig. 1E).^47^ This object-aware approach represents an advance upon pixel-level memory alone,^49,50^ as object awareness mitigates the propagation of pixel-level error, particularly during occlusions, and supports the stability of object detection over time.

Cutie performs semi-supervised VOS by initiating directional inference using the first frame’s target object segmentation as a starting point. We piped pixel-prompted vole segmentation masks generated by Segment Anything Model^51^ into Cutie’s inference engine (Supp. Fig. 2A). We chose to split two-hour-long behavior videos into approximately five-minute-long clips, with each clip bookended by experimenter-vetted, segmented keyframes (Fig. 2A).

**Figure 2:**
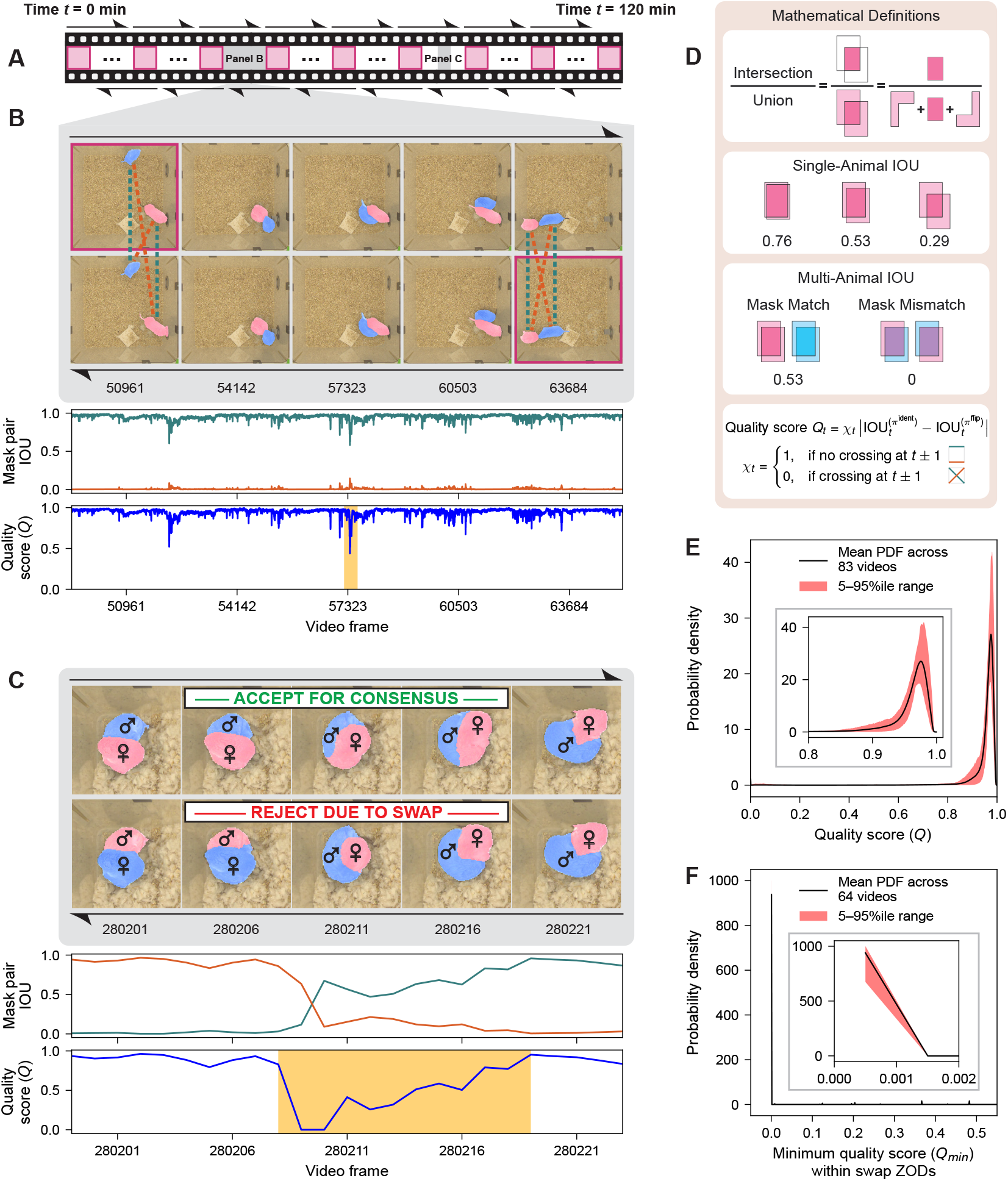
Bidirectional video object segmentation (VOS) combined with mask intersection over union (IOU) enables precise detection of segmentation errors and supports robust consensus mask generation. A. Schematic of a two-hour behavior video (not to scale). User-validated keyframe masks (magenta-outlined squares) initiate Cutie-based segmentation in both forward and reverse directions (black hemi-arrows). Gray-shaded segments are shown in expanded temporal detail in B and C. B. Example segment (gray region from A) illustrating bidirectional inference. Forward masks propagate from the upper left and reverse masks from the lower right. Identity mapping comparisons (same-animal masks across directions; teal dotted lines) yield high IOU π^ident^ values (teal curve), whereas flipped mapping comparisons (different-animal masks; burnt orange dotted lines) produce low IOU π^flip^ values (burnt orange curve). Quality score is defined as the absolute difference between IOU π^ident^ and IOU π^flip^, with a score of 0 assigned near curve intersections. Low-quality regions (score < 0.5; butterscotch shading) are flagged for manual review. C. Example zone of disagreement (ZOD, butterscotch shading) resulting from an identity swap during reverse inference. Ground truth identities are denoted by male and female symbols. Because the reverse direction contains the swap, forward masks are retained for consensus mask generation. IOU π^ident^ and IOU π^flip^ curves cross in an “X”-shaped pattern typical of one-direction swaps. D. Schematic definition of IOU and quality score calculations used for bidirectional mask comparison. Single-animal examples illustrate how even small mask displacements can cause a marked reduction in IOU, with values that remain below the norm observed in our dataset (see E). Multi-animal examples demonstrate that IOU reflects both mask area overlap and correct identity assignment. E. Mean probability density function (PDF) of framewise quality scores across 83 two-hour videos. The red shading indicates the 5^th^-95^th^ percentile range. Most frames exceed a score of 0.9, reflecting high bidirectional agreement (see benchmark examples in Fig. 2D). Inset provides a magnified view of the distribution in the rightmost region. F. Mean PDF of the minimum quality scores for all ZODs that contain a swap in one or both directions. The red shading indicates the 5^th^-95^th^ percentile range across 83 videos. These ZODs exhibit markedly lower minimum scores compared to the overall distribution. Inset provides a magnified view of the distribution in the leftmost region.

This choice was motivated by the following goals: (1) to accelerate inference with highly parallelized, cluster-based computing, (2) to reduce GPU memory requirements per clip, and (3) to balance the tracking advantages of memory with the long-term risk of error propagation. Next, starting from each boundary keyframe, we performed bidirectional VOS, yielding two sets of segmentation masks for each video clip (Fig. 2A-C). Since forward and reverse inference for a given video clip proceed from distinct segmentation prompts and traverse memory space in different ways, the two sets of masks are effectively independent.

To assess segmentation fidelity, we compared forward and reverse Cutie-generated masks using framewise intersection over union (IOU; Fig. 2D). We reasoned that accurate, identity-consistent masks for the same animal should yield IOU values approaching 1.0 (Fig. 2B,C; teal dotted lines and IOU π^ident^ curves), while comparisons between masks corresponding to different animals should yield IOU values near 0.0 (Fig. 2B,C; burnt orange dotted lines and IOU π^flip^ curves). We defined quality score as the absolute difference between the IOU values of the two permutated mappings between forward and reverse masks, π^ident^ and π^flip^, with frames flanking each intersection point between the two curves assigned a score of 0 (Fig. 2D). This scoring scheme assigns higher values to frames with strong bidirectional agreement and lower values to those more likely to contain identity mismatches. In aggregate, quality scores were consistently high across full videos, reflecting strong agreement between forward and reverse segmentations and highlighting the rarity of ambiguous regions (Fig. 2E). Since segmentation errors are typically history-dependent, we reasoned that they would not occur on the same frame and in the same way in both directions. Thus, regions where the two segmentations diverged – termed zones of disagreement (ZOD) – were likely to reflect one of three error modes: a forward-specific error, a reverse-specific error, or a bidirectional failure. In practice, mask identity swaps occurring in only one direction produce a characteristic “X”-shaped intersection between IOU π^ident^ and IOU π^flip^ curves, signaling a candidate ZOD for review (Fig. 2C).

To support efficient ZOD review, we developed a graphical user interface that displays framewise quality scores and highlights ZODs for annotation (Supp. Fig. 2B). Each ZOD is summarized by its lowest-scoring frame, enabling users to prioritize review of the most error-prone ZODs. In most cases, users simply select the preferred segmentation direction (forward or reverse) based on visual inspection, correcting mask identity swaps with minimal effort. When applied across 83 two-hour videos of dyadic interactions between voles, this process yielded exceptionally low error rates: a one-direction swap rate of 1.01 ± 0.14 per 100,000 frames and a both-direction swap rate of just 0.21 ± 0.05 per 100,000 frames (right-hand side of Fig. 3A). In cases where a swap occurs in only one direction, the opposite direction can be used to generate identity-validated consensus segmentations (see example in Fig. 2C).

**Figure 3:**
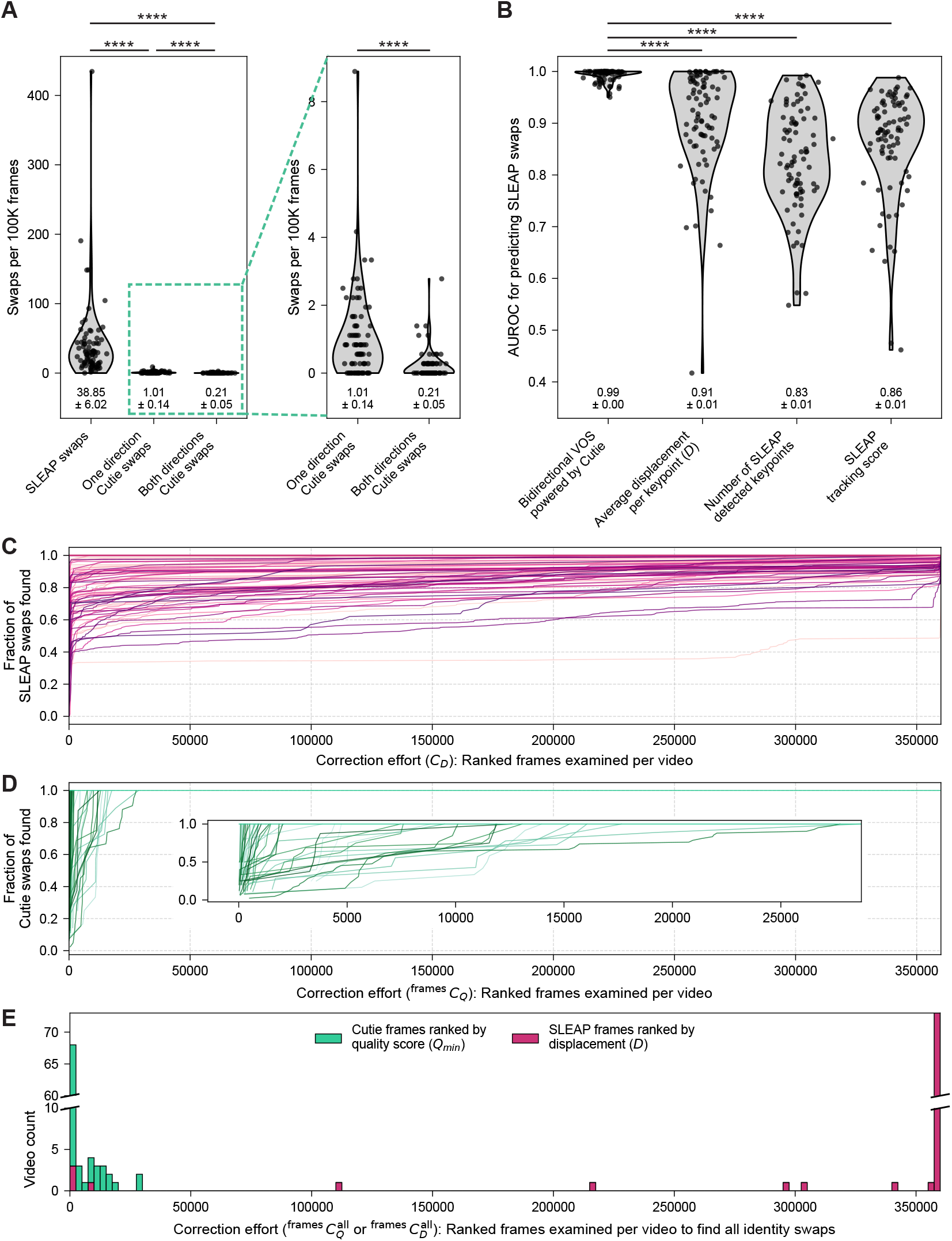
Cutie-derived consensus masks enable high-accuracy identity correction with minimal user effort. A. Identity swap rates for SLEAP keypoints with post hoc tracking versus Cutie-based bidirectional segmentation across 83 two-hour videos (360,000 frames each). Violin plots show the distribution of swaps per 100,000 frames; individual videos are shown as dots, and means ± s.e.m. are indicated below. Mann-Whitney U tests revealed highly significant differences between Cutie and SLEAP methods (****, *p* < 0.0001). Due to the disparity in scale, Cutie swap rates are also shown on an expanded y-axis. Most Cutie swaps affect only one direction of inference, enabling frames from the other, error-free direction to be used for consensus mask generation. B. Area under the ROC curve (AUROC) values for swap detection using Cutie-derived consensus masks versus four SLEAP-based heuristics: average motion energy per keypoint, SLEAP tracking score, and number of SLEAP detected keypoints. Consensus masks consistently outperformed all SLEAP-based metrics, yielding higher mean AUROC and lower variability across videos. C. Fraction of SLEAP swaps recovered as a function of correction effort C_E_, for which adjacent frame pairs were ordered by descending motion energy. Each curve represents a single two-hour video (360,000 frames). D. Fraction of Cutie swaps recovered as a function of correction effort 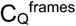, for which ZODs are ordered by ascending quality score. Each curve represents a two-hour video. Inset shows an expanded view of the leftmost portion of the curve. E. Number of frames manually reviewed per video to recover all identity swaps using either Cutie ZOD flagging or SLEAP keypoint motion energy; equivalent to the correction effort required to achieve 100% swap discovery. Histogram shows the distribution across all 83 videos. Cutie required substantially less manual inspection, with a median of 411 frames reviewed per video (out of 360,000), compared to the near-complete review needed for SLEAP-based methods.

We simulated worst-case identity swaps to test the limits of our approach. Specifically, we introduced synchronous or near-synchronous swaps in both inference directions (Supp. Fig. 4C-E). In these cases, swap detection hinges on the degree of disagreement between segmentation streams. For example, when swaps are offset by just one frame, the corresponding IOU (Supp. Fig. 4D) drops sharply to 0, well below the proposed ZOD threshold and thus easily flagged. However, when swaps occur at the same frame in both directions, the drop in IOU is less drastic (Supp. Fig. 4E) or even nonexistent (Supp. Fig. 4C). In practice, bidirectional swaps did not occur simultaneously and with identical segmentation errors in our large two-animal tracking dataset; the independence of forward and reverse inference led to divergent error timing and propagation, enabling reliable detection.

**Figure 4:**
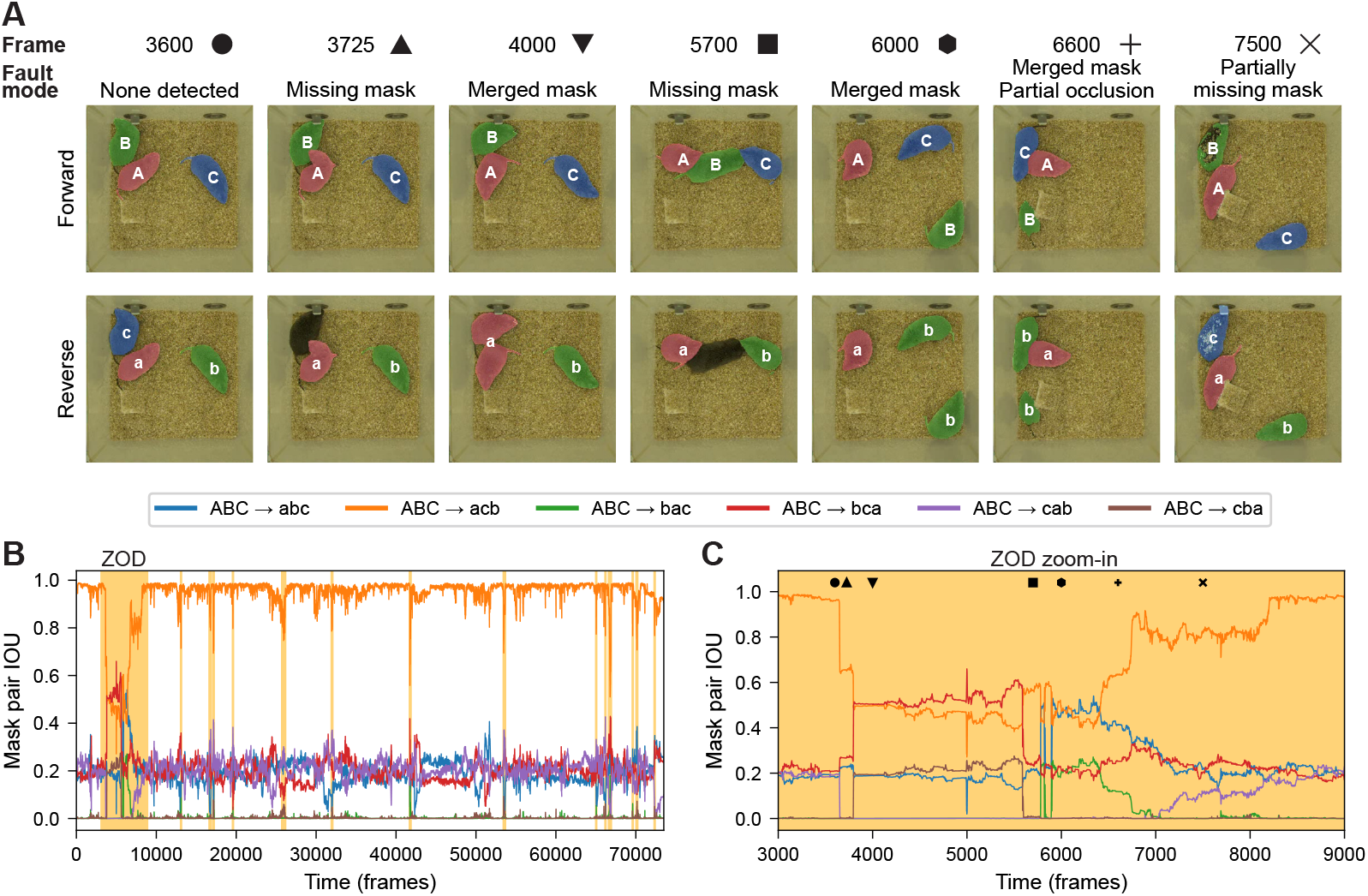
Proof-of-principle for three-animal tracking with bidirectional VOS. A. Forward (uppercase letters) and reverse (lowercase letters) segmentation masks overlaid on raw video frames, illustrating representative segmentation fault modes. Frame 3600 shows consistent segmentation in both directions; all other frames depict faults in one direction or the other. Symbols mark each frame (corresponding IOU values in panel C). B. Pairwise IOUs for all six possible mask-to-mask mappings (see legend) over a 24.5-minute recording at 50 fps. Intervals with reduced dominant IOU are highlighted with butterscotch shading. The color legend is displayed in the box above. C. Expanded view of the large ZOD from panel B. Symbols indicate the frames shown in panel A, linking each segmentation fault mode to its corresponding IOU values.

### Bidirectional video object segmentation significantly outperforms pose estimation with post hoc tracking in identity preservation

To correct SLEAP identity errors, we computationally aligned keypoints to Cutie-derived consensus segmentation masks (see flow chart in Supp. Fig. 3D). This was done by computing an accuracy-weighted fraction of keypoints that best matched the masks in each direction (see Methods). We confirmed that this procedure correctly resolved all of the SLEAP swaps in the manually curated ground truth SLEAP dataset (Supp. Fig. 1A, 1B). As expected, bidirectional Cutie dramatically reduced identity swaps compared to SLEAP, yielding approximately 185-fold fewer swaps (0.21 ± 0.05 versus 38.85 ± 6.02 swaps per 100,000 frames; left-hand side of Fig. 3A, Supp. Fig. 3C). These consensus masks exhibited markedly higher discriminative ability for detecting SLEAP identity swaps than any of the three post hoc SLEAP-based heuristics, which showed both lower mean AUROC values and greater variability across videos (Fig. 3B).

In large-scale behavioral video analysis, minimizing the manual effort needed to identify and correct identity swaps – specifically, the number of frames requiring review – is as important as classification performance itself. SLEAP-based heuristics often require manual inspection of nearly every frame to capture all swaps (Supp. Fig. 1D; Fig. 3C, 3E). In contrast, our Cutie-based pipeline enables a more targeted approach: by flagging only ZODs for review, the annotation burden is substantially reduced (Fig. 3D, 3E). In most two-hour-long videos (360,000 frames), fewer than 10 ZODs and fewer than 1,000 frames require inspection (Fig. 3E; Supp. Fig. 3A, 3B). Across the full dataset of nearly 30 million frames, identity swaps occurred almost exclusively in ZODs with quality scores near 0.0, and none were observed above a score of 0.5 (Fig. 2F; Supp. Fig. 3A, 3B). Reviewing all ZODs up to this criterion threshold reliably captured all swaps. In contrast, there does not exist an equivalent criterion threshold for SLEAP-based heuristics, which show no clear cutoff that reliably distinguishes swaps from non-swaps.

### Bidirectional video object segmentation extends to the tracking of three interacting animals

Beyond dyadic interactions that comprise aspects of pair bonding, complex social dynamics evolve within groups of animals. We therefore tested whether our bidirectional VOS approach generalizes to tracking three unmarked animals. First, we describe how the core computational framework scales from two to more individuals. Second, we consider edge cases of identity swaps that risk evading detection by ZOD-based review. Lastly, we offer general guidance for adapting this strategy to even larger groups.

At the heart of our approach is the comparison of forward and reverse segmentations via pairwise IOU values. For two animals, only two mask permutations exist, yielding IOU π^ident^ and IOU π^flip^. For three animals, this extends to all 3! = 6 possible permutations. Within a well-segmented interval, one mapping should yield a complete identity match (3/3 animals), with IOU values near 1.0 (Fig. 4B, C, Supp. Fig. 5A). Three mappings will yield only a partial identity match (1/3 animals), and the remaining two will yield no identity match (0/3 animals). For example, in Fig. 4B and Supp. Fig. 5A, the ABC → acb mapping (i.e., forward labels A-B-C matching reverse labels a-c-b) shows a clear IOU distribution centered near 1.0, indicating identity correspondence.

The next step is to use IOU values to identify candidate ZODs for manual review, with the goal of capturing all potential identity swaps. Unlike the two-animal case, where a quality score *Q* was computed, here we adopt a more general approach: for each frame, we determine the dominant mapping as the one with the highest IOU among the six. A change in dominance – i.e., when IOU curves intersect – can indicate a potential identity swap in one inference direction but not the other, and the surrounding interval is flagged as a ZOD. Based on the distribution of IOU values across mappings (Supp. Fig. 5A) as well as the IOU values of known segmentation fault modes (Fig. 4A, C), we find that reviewing all regions where the dominant IOU falls below 0.8 (approximately 5.3% of frames in the example video; Supp. Fig. 5B) is sufficient to identify all potential swaps. In this example, no identity swaps were present in either inference direction, although the reverse segmentation showed several “merged mask” faults (Fig. 4A, C), where one mask temporarily encompassed two animals and another mask disappeared.

While tracking three animals introduces a larger set of potential bidirectional swap configurations (e.g., forward swaps between animals 1 and 2 concurrent with reverse swaps between animals 2 and 3 within a ZOD), most such cases can be readily detected from IOU curve crossings. The critical edge cases arise when symmetric swaps occur simultaneously in both directions between the same pair of animals. These synchronous dyadic swaps represent the natural three-animal extension of the two-animal failure modes illustrated in Supp. Fig. 4C-E, and they pose the greatest challenge for maintaining stable identity assignments. However, as in the two-animal case, such perfectly synchronous symmetric swaps did not arise in practice.

As the number of tracked animals *n* increases, each individual contributes a smaller fraction of the total overlap (approximately 1/*n*, assuming similar mask sizes). Consequently, the examination threshold for the dominant IOU should be adjusted upward to maintain sensitivity. Furthermore, during manual annotation of identity swaps within ZODs, users must not only confirm the presence of a swap but also specify the new identity permutation – that is, the updated mapping of masks to animal identities resulting from the swap.

## DISCUSSION

We present a novel approach for achieving high-confidence, identity error-free tracking of markerless, interacting animals using bidirectional video object segmentation (VOS). Our pipeline integrates four key steps: (1) user-guided generation of keyframe segmentation masks, (2) bidirectional VOS inference initiated from those keyframes, (3) user-supervised resolution of zones of disagreement (ZODs), and (4) automated alignment of pose estimation keypoints to identity-validated segmentation masks. Together, these components enable long-term, multi-animal identity tracking with drastically reduced manual annotation effort. For two interacting animals, our method requires several orders of magnitude less user intervention than direct correction of identity swaps in pose estimation data. While standard unidirectional VOS already produces fewer identity errors than pose estimation with post hoc tracking, bidirectional inference is a powerful tool for detecting and resolving the rare errors that persist.

Unlike our approach, traditional identity-tracking strategies rely on physical markers such as dyes,^52,53^ bleach,^10,54^ fur shaving,^35,55,56^ or RFID tags^35,57^ – all of which require additional handling that can be stress-inducing for animals.^36^ Visual markings can alter appearance in ways that disrupt natural behavior, particularly in species sensitive to visual social cues. Similarly, chemical agents like paint or bleach may introduce olfactory signals that interfere with behavior in chemosensory-reliant animals. Implanted RFID tags avoid these sensory confounds but suffer from poor spatial and temporal resolution, signal collision, and detection dropout, especially in dense social contexts.^37–40^ Moreover, RFID systems lack pose information and are not readily compatible with keypoint-based tracking. By contrast, our method achieves identity-resolved tracking with minimal behavioral disruption and high temporal and spatial precision, while remaining compatible with both pose estimation data and traditional physical markers.

These advantages open the door to several powerful applications. We foresee three primary use cases for our bidirectional VOS approach: (1) studying behavioral change across extended timescales, (2) analyzing social dynamics in multi-animal environments, and (3) achieving accurate identity tracking with limited effort when pose estimation is not required.

First, our method facilitates long-term, continuous tracking of social interactions in naturalistic settings, making it especially well-suited for studying dynamic, context-dependent, and low-frequency behaviors that unfold over hours, days, or even weeks. These include patterns that fluctuate with circadian rhythms,^58^ shift as a result of prolonged social experience,^31^ or evolve gradually during development.^59,60^ For example, our approach can track the emergence and maintenance of pair bonds, the evolution of social behaviors following isolation, and gradual shifts that reflect slow-changing internal states. While brief video samples remain useful for many experimental paradigms, continuous tracking opens the door to understanding not only *what* behaviors are expressed, but *when* and *how* they evolve. This capability is particularly valuable in behavioral neuroscience, where there is increasing emphasis on studying naturalistic behaviors across larger cohorts. As experiments grow in scale, such as with high-throughput genetic or pharmacologic screens,^61^ methods that reduce tracking burden without compromising accuracy will be critical.

Second, our method enables accurate, identity-preserving tracking in group-housed or otherwise socially complex environments. Many forms of social behaviors, such as selective affiliation,^35,62^ social buffering,^63,64^ or the emergence of dominance structures,^65–67^ cannot be easily captured through isolated or pairwise encounters. They depend on the presence of multiple individuals, the evolving relationships among them, and the full context of group interactions. Our approach should allow researchers to continuously track all individuals in a group, even during dense physical interactions or periods of social reorganization. This makes it possible to ask how group structure emerges and stabilizes, how individuals adapt their behavior in response to others, and how social information spreads across a network. These questions are central to ethologically grounded studies of social learning, in-group versus out-group discrimination, and collective behavior in mammals and other socially organized species.

Third, for users primarily interested in tracking individual identities in social contexts – without the added complexity of pose estimation – our method offers a high-accuracy solution that is immediately applicable. Because identity is derived directly from segmentation mask continuity, our approach avoids the need for training or fine-tuning a pose estimation model, which typically requires a large set of manually curated keypoint labels. In contrast, pose estimation pipelines often rely on supervised learning and benefit from dataset-specific annotation, making them more labor-intensive for users whose primary goal is reliable identity tracking.

It is important to consider several limitations of our bidirectional VOS approach. First, complete occlusion of visually identical animals – such as when they disappear beneath objects in the environment – can result in identity loss, especially if all animals vanish simultaneously under the same obstruction for an extended period. In such cases, identity continuity cannot be reliably maintained through segmentation alone and may instead require physical identifiers that distinguish one animal from another. Second, there is likely a practical upper bound on the number of animals that can be tracked with high confidence using bidirectional VOS, due to a combination of behavioral and computational constraints. As group size increases, animals are more likely to engage in complex interactions – such as prolonged huddling which results in the occlusion of more than one animal at a time – that challenge segmentation-based identity resolution.

While our default ZOD resolution strategy prioritizes identity continuity, users interested in improving keypoint localization accuracy may opt to resolve ZODs using a more stringent quality score threshold. This approach yields more refined consensus masks, which could in turn be used to prune keypoints that fail to align with high-confidence segmentation boundaries.

Beyond using segmentation masks as a post hoc filter for pose data, we envision future architectures in which shape information from segmentation is incorporated as a prior during pose estimation itself, potentially improving both keypoint accuracy and mask quality in a unified inference process. Toward this goal, we propose the development of integrated models that jointly perform video object segmentation and pose estimation, allowing mutual constraints between tasks. In parallel, we suggest exploring architectures that permit message-passing between forward and reverse segmentation passes, which could substantially reduce the need for post hoc bidirectional comparison by resolving mask identity conflicts during inference.

## Materials and Methods

### 1 Animal Rearing

All animal care and procedures were approved by the Institutional Animal Care and Use Committee at the University of California, San Francisco (UCSF). A total of 186 sexually naïve adult prairie voles (*Microtus ochrogaster*) and meadow voles (*Microtus pennsylvanicus*), 7–9 weeks of age, were used in this study. Both males and females were included, with sex determined by gonadal inspection at weaning.

Voles were bred in our laboratory from systematically outbred wild-caught stocks (prairie voles: near Champaign, IL; meadow voles: Hampshire County, MA) and maintained in our UCSF facilities. *Oxtr*–/– prairie voles were derived from a previously generated line, and wild-type controls were obtained from the same outcrossing background line. Breeding pairs were housed in large plastic cages (10.5 in × 19 in × 8 in; Ancare, R20 Rat/Guinea Pig) on Paperchip bedding (Shepherd Specialty Papers).

Weaned voles (21–25 days) were group-housed in clear plastic cages (45 cm × 25 cm × 15 cm; Innovive, Innocage IVC Rat) with Paperchip bedding, then transferred to our lab housing facility and re-housed on Sani-Chips woodchip bedding (P.J. Murphy, Forest Products Corp.). Each group cage contained 2–6 same-sex siblings or age-matched weanlings, two cotton nestlets, and a large PVC elbow tube. Food and water were provided *ad libitum*.

For pair housing, voles were kept in 30.8 cm × 30.8 cm × 18.7 cm cages (Thoren, Maxi-Miser Model #4) on Sani-Chips bedding with two cotton nestlets and two small plastic tubes. All animals were maintained on a 14:10 light–dark cycle.

### 2 Behavior

#### Assays

We recorded social behavior videos of the first meeting between sexually naive, opposite-sex animals. On the day of an assay, voles were transferred from the housing room to the procedure room at least half an hour prior to testing. Voles were briefly restrained by the experimenter to confirm identity. Two age-matched, opposite-sex voles were placed into a clean cage, one after the other, and behavior was recorded for two hours. During all assays, all tubes, food hoppers, and water bottles were removed from the cage. Importantly, animals were completely markerless—they were not subject to fur clipping, fur dyeing, fur bleaching, or radio-frequency identification (RFID) tagging prior to assay initiation.

#### Video Acquisition and Encoding

We employed four Basler ace acA2040-55uc cameras (3MP resolution) for simultaneous video recording, capturing 2048 × 1536 pixels at 50 frames per second with Bayer RGGB8 encoding via a dedicated PCI Express USB 3.0 interface card. Multi-camera video compression was conducted in real-time on a workstation with a consumer graphics card. We used `campy`^1^ to generate `.mp4` video with `ffmpeg` and `libx264` at Constant Rate Factor 26. All videos captured slightly exceeded 2 hours in duration to accommodate experimental setup and animal identification procedures. Videos were later trimmed to the first usable frame after the addition of a second vole, and analyses all terminate at frame 360,000 (*t* = 120 minutes) in each video.

### 3 Pose Estimation

#### SLEAP Inference

We trained a SLEAP model to generate pose estimation data on pairs of voles.2 Our training set included 2,423 frames of prairie and meadow voles in the same home cage enclosures used in the rest of our videos. We used a bottom-up network with a node sigma of 2.5 and edge sigma of 100. The following SLEAP augmentations were applied: rotation range [−180^∘^, 180^∘^], scale [0.7, 1.3], Gaussian noise [5.0, 1.0], contrast [0.5, 2.0], brightness [0, 10], and horizontal flips. With these parameters, we achieved a training validation loss of 0.000423.

The SLEAP skeleton configuration included keypoints for the nose, left and right ears, three along the spine, left and right flanks, rump, tail base, and tail tip. Due to frequent occlusion of the tail beneath the animal’s body or bedding, the tail base and tail tip were inconsistently detected. As a result, these two keypoints were excluded from downstream analyses of average keypoint displacement (*D*) and alignment to segmentation masks (Supp. Fig. 1C, Fig. 3A).

#### SLEAP Tracking

We employed several pose tracking methods from SLEAP version 1.3.0, parameterized as follows:

| tracker | Simple | Simple | Flow |
| --- | --- | --- | --- |
| target_instance_count | 2 | 2 | 2 |
| pre_cull_to_target | True | False | 2 |
| similarity | Instance | Instance | Instance |
| match | Hungarian | Hungarian | Hungarian |
| track_window | 5 | 5 | 5 |
| min_new_track_points | 3 | 3 | 3 |
| min_match_points | 3 | 3 | 3 |

*\*Additional parameters relevant for the flow tracker method:*
`img_scale = 0.5, of_window_size = 21, of_max_levels = 3`

The SLEAP tracking process sometimes produced more than two tracks, often consisting of partial trajectories, despite specifying only two animals. In some cases, one partial track terminated at the same frame another began. We applied SLEAP’s track-cleaning tool to merge such pairs of partial tracks, aiming to obtain two continuous tracks spanning the entire video. When the track-cleaning tool failed to reduce the output to exactly two tracks, we implemented custom code to enforce the generation of two complete trajectories. Identity swaps introduced by the cleaning process were negligible compared to those generated during tracking.

#### Tracking Metrics

##### Displacement

To quantify frame-by-frame movement in each animal, we computed a displacement metric based on SLEAP keypoints. The motivation for this analysis was to obtain a compact, interpretable summary of whole-body motion that could be compared across time, individuals, and behavioral conditions. For example, if an animal remains completely still between two adjacent frames and keypoints are consistently detected, its displacement will be zero. In contrast, if the animal makes a rapid movement—such as leaping halfway across the cage—the displacement will be high.

We defined a displacement metric *D* computed per animal and per frame. For each animal *a* ∈ {1, 2} and each frame pair (*t, t* + 1) with *t* ∈ {1, 2, …, *T* − 1}, where *T* denotes the total number of frames, we identified all keypoints that were valid (non-NaN and non-negative) in both frames. Denoting the number of shared valid keypoints as 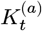, we defined the average displacement per keypoint as:

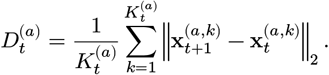

Here, 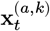 ∈ ℝ^2^ is the 2D coordinate of keypoint *k* for animal *a* at frame *f*, and ‖*⋅* ‖ denotes the Euclidean norm.

Importantly, this metric is sensitive to tracking errors: if two animals are stationary but their identities are swapped between frames, the resulting displacement may be spuriously elevated due to the mismatch in body part positions. However, displacement can also fail to detect true identity swaps—for instance, when two animals are in close proximity and keypoint detections are sparse or inaccurate, a swap may produce little apparent displacement.

For each video, we ranked all adjacent frame pairs in descending order of displacement *D* to guide the discovery and correction of identity swaps. The cumulative fraction of known SLEAP identity swaps discovered at a given correction effort is denoted by *SD*(*CD*), and computed as:

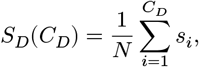

where:

1. *CD* is the number of top-ranked frame pairs examined (i.e., the correction effort),
2. *si* ∈ {0, 1} indicates whether frame pair *i* contains a swap (1 = swap, 0 = no swap),
3. 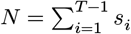is the total number of known identity swaps in the video, across *T* − 1 adjacent frame pairs.

Plotting *SD*(*CD*) as a function of *CD* yields an interpretable curve: steep initial rises indicate that high-displacement frames are enriched for identity swaps and are therefore efficient to prioritize for manual review, while subsequent flattening of the curve reflects a long tail of sporadic identity swaps that occur even in frames with low displacement (Supp. Fig. 1Di).

##### SLEAP tracking score

We recorded the instance-level track matching score outputted by SLEAP tracking to assess its sensitivity and specificity for predicting SLEAP identity swaps.2

##### SLEAP number of keypoints detected

Since tracking errors are partially attributable to missing or erroneously detected keypoints, we also recorded the number of keypoints (body parts) detected per animal 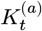.

### 4 Video Object Segmentation (VOS)

#### Cutie Overview

We use Cutie for semi-supervised video object segmentation (VOS),3 initializing segmentation with a single keyframe (a video frame annotated with ground-truth masks for the two voles). In our initial trials, we found that Cutie produced stable, high-quality segmentations for our videos over intervals exceeding 20,000 frames when initialized with ground-truth masks from the first frame of a video clip.

Cutie’s baseline performance was strong, likely due to its incorporation of both object-level and pixel-level memory. Nonetheless, we observed occasional segmentation errors, including partial mask loss, erroneous inclusion of environmental features (e.g., bedding or cage ports), and—in rare instances—identity confusion between animals. Because Cutie’s inference is history-dependent, we reasoned that error patterns might vary depending on where inference begins. We therefore implemented a bidirectional inference strategy, running Cutie forward and backward in time for each video from distinct keyframes with ground-truth masks (the first and last frames of each clip). This yielded two independent segmentation streams per frame, increasing robustness and enabling systematic detection of inconsistencies, since it is unlikely that forward and reverse inferences would exhibit identical pixel-level errors over the same interval.

Cutie inference was performed with a code snapshot from June 4, 2024 (GitHub commit hash `e869fab5e653b118c475cbafe0b2322b37a7e320`). We used the `cutie-base-mega.pth` weights (SHA256 hash `9c05402ee36d3a356fb72715d263ba7e1ea06ad3bada48c1306491792da43023`) for inference.

#### Cutie Segmentation Clip Interval

Although Cutie reliably segmented video clips exceeding 20,000 frames in length, we chose to perform inference on clip lengths of 10,000–20,000 frames for several reasons. First, we experienced much higher availability of GPU computing resources on the UCSF cluster when compute jobs were constrained to complete in under two hours, which was more reliably achieved with shorter segment lengths. Second, our experiments often include periods in which voles are engaged in prolonged bouts of close-range behaviors. Our validation of Cutie segmentations suggested an advantage to selecting keyframes where voles were physically separated from each other. These constraints contributed to variability in segment lengths. Across 83 introduction videos, we observed a mean segment length of 13,825 frames (median 14,789 frames).

For the three-animal video recording, bidirectional Cutie inference was performed for a continuous 75,000-frame segment.

#### Keyframe Generation

To streamline keyframe generation, we developed a Python-based annotation tool (Supp. Fig. 2A) powered by Meta AI’s Segment Anything Model (SAM).4 Because SAM inference in CPU mode was prohibitively slow, we implemented a client–server design: a lightweight client application that could be run on standard laptops, and a server component packaged in a Docker image and deployed on short-term GPU instances rented through runpod.io. This configuration enabled parallel labeling by multiple researchers at a total compute cost of under $200, including development and testing. To identify keyframes for VOS, we first sampled approximately 250 candidate frames per video, always including the first and last usable frames from each 360,000+ frame recording. From this set, we prioritized frames that met the following criteria: (1) first and last frames of the video, (2) animals clearly separated or, when not possible, minimally occluded with a visible boundary between them, and (3) inter-keyframe intervals of less than 20,000 frames.

For annotation, users placed one or more positive prompts on each vole and, when needed, negative prompts to exclude background elements. SAM inference then yielded candidate segmentation masks, from which users selected the best pair corresponding to the two voles. The chosen masks were merged into a single keyframe mask image.

To refine annotations, we developed a complementary proofing tool that automatically removed small spurious artifacts occasionally produced by SAM and allowed manual mask adjustment in cases where SAM segmentation was suboptimal.

#### Assembly of Segments Across Videos

Adjacent Cutie inference segments were configured to overlap by a single frame—namely, the boundary keyframe containing user-defined ground-truth masks used to seed bidirectional semi-supervised VOS. To generate a continuous sequence of forward or reverse masks, we evaluated identity alignment across segment boundaries by comparing overlapping masks. Specifically, for each pair of overlapping frames, we computed the sum of pixelwise intersection for both identity-aligned and identity-flipped configurations. The configuration yielding the greater intersection value was used to determine the correct stitching orientation. Across our dataset, this procedure consistently produced clear, unambiguous stitching outcomes.

We then downsampled the resulting frame sequences by a factor of four in both height and width dimensions (from 1536 × 2048 to 384 × 512) to facilitate data handling and reduce storage requirements. This process generated compressed, random-access .h5 files for each video.

### 5 Mask Generation for *n*-Animal Tracking

#### Permutational Comparison of Bidirectional Masks

To evaluate the consistency of bidirectional VOS, we compare forward and reverse segmentation masks by computing intersection-over-union (IOU) values across all possible identity permutations.

Let 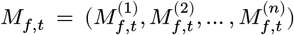and 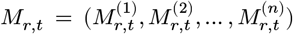 denote the forward and reverse segmentation masks at frame *t* for *n* animals. Each 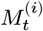 is a binary mask of the same spatial resolution indicating the pixels belonging to animal *i*.

However, since forward and reverse inference for a given video segment are initialized from distinct keyframes, there is no guarantee that corresponding animals are marked by the same mask labels within *M*_*f*_ and *M*_*r*_. Let *𝒫*_*n*_ denote the set of all *n*! permutations of the index set {1, 2, …, *n*}. For each permutation *π* ∈ *𝒫*_*n*_, we define a permuted version of the reverse masks:

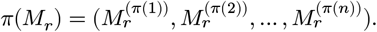

For each permutation *π*, we compute the combined IOU as:

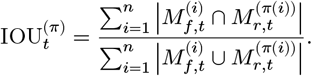

This formula sums the intersection areas across all aligned mask pairs and divides by the sum of their unions, yielding a single scalar that quantifies the overall overlap quality under permutation *π*.

The permutation 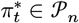 that maximizes IOU^(*π*)^ at time *t* is considered the most dominant alignment between forward and reverse masks:

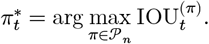

The corresponding maximal value, 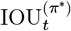, serves as a framewise score that quantifies segmentation consistency under the best identity mapping. These scores are subsequently used to flag candidate ZODs for manual review.

#### Detection of Candidate ZODs via Dominant IOU Transitions and Thresholding

To identify candidate *zones of disagreement* (ZODs) for manual review, we implemented a two-phase procedure based on the behavior of the dominant IOU time series, 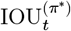, across time.

##### Phase 1: Dominance Transitions via IOU Intersections

We detect potential identity inversions by identifying timepoints where the dominant permutation 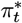 changes:

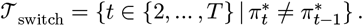

Around each *t* ∈ 𝒯switch, we define a temporal window of radius *r*. The set of candidate ZODs from this phase is thus

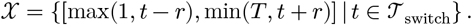

Overlapping or adjacent intervals in 𝒵 are merged to form a consolidated set of candidate regions.

##### Phase 2: Threshold-Based Detection of Low-Confidence Regions

To identify bidirectional swaps that may not yield a change in 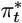, we apply a threshold to the dominant IOU time series itself:

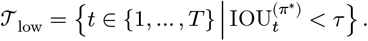

Here, τ ∈ (0, 1) is a user-defined threshold selected based on empirical performance based on the distribution of IOU values and known segmentation fault modes. For *n* = 3, a threshold of 0.8 should be sufficient for capturing all potential identity swaps based on our data. We segment 𝒯low into contiguous intervals:

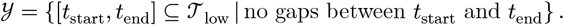

Each interval defines a candidate ZOD, capturing regions where segmentation agreement is consistently poor, even in the absence of switches in permutation dominance.

##### Final Candidate ZOD Set

The final set of candidate ZODs is obtained by merging the intervals detected in Phase 1 and Phase 2:

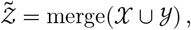

where merge(*⋅*) denotes the consolidation of overlapping or adjacent intervals into non-overlapping candidate regions. This procedure ensures comprehensive coverage of both unidirectional and bidirectional inconsistencies while minimizing redundant manual review.

#### Manual Annotation of ZODs and Consensus Mask Assembly

Once candidate ZODs are identified through automated detection, yielding the set 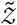, the next step is manual refinement of their temporal boundaries. This produces the set 𝒵, in which each 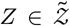 has been adjusted, if necessary, so that it begins and ends on frames with high-quality segmentations.

Subsequent annotation determines whether each *Z* ∈ 𝒵 contains an identity swap. If a swap is detected, the annotator specifies the corrected identity mapping by providing an updated permutation *π*_corrected_ ∈ *𝒫*_*n*_, where *𝒫*_*n*_ denotes the set of all possible permutations of the *n* animal identities. This permutation defines the proper assignment of instance masks (e.g., A, B, C) to animal identities (e.g., 1, 2, 3) for the affected frames.

Directional masks in which a swap has been annotated are designated as “garbage” and discarded, while the other direction is retained for consensus mask assembly. If both forward and reverse outputs are annotated as containing swaps, the ZOD is labeled as a double-garbage ZOD (dgZOD), and no output is retained. When segmentation errors occur in temporally separated portions within a ZOD, annotators may subdivide it into smaller subregions, each with its own consistent corrected mapping *π*_corrected_.

Through this process, manual annotations determine whether to retain the forward direction, retain the reverse direction, or discard both. For non-ZOD regions, either direction may be selected interchangeably, as their outputs are nearly equivalent. Concatenating the selected masks across ZODs and non-ZODs produces a single stream of consensus masks that are free of identity swaps and exhibit improved segmentation quality.

#### Two-Animal Tracking

##### Specialized Definitions

Although the dominant-permutation framework generalizes naturally to *n* ≥ 3 animals, the case of *n* = 2 admits a simpler formulation. Because there are only two possible permutations, one can bypass the dominant-permutation definition and instead work directly with the IOUs under these mappings:

- **Identity mapping** *π*^ident^: *π*^ident^(1) = 1, *π*^ident^(2) = 2
- **Flipped mapping** *π*^flip^: *π*^flip^(1) = 2, *π*^flip^(2) = 1

We denote the corresponding IOU values as 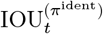 and 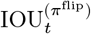. The framewise quality score is then defined as:

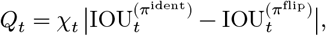

where

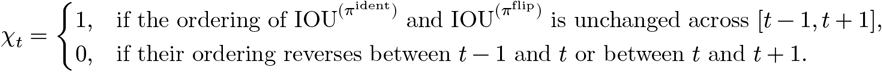

Here, the indicator χ*t* acts as a flip-aware suppression term, setting the score to zero in the vicinity of IOU crossings.

This quality score takes values near 1 when the paired masks agree on animal extents regardless of identity assignment, and values near 0 when they diverge. It therefore enables detection of ZODs, including identity inversions (𝒳) and segmentation disagreements (𝒴), across paired Cutie inference streams. Such zones can only be identified when forward and reverse inferences do not fail in precisely the same way (i.e., with matching pixel-level error patterns) over the same interval (see Supp. Fig. 4).

##### Quantitation of ZOD Annotation Effort

For each *Z* ∈ 𝒵, we computed a summary statistic based on framewise quality scores *Qt*. Formally, we defined

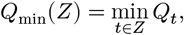

where *t* ∈ *Z* denotes the set of frame indices spanned by ZOD *Z*. Intuitively, *Q*_min_(*Z*) captures the most severe segmentation disagreement within the region and was used to rank ZODs by priority for manual inspection. In all subsequent calculations, ZODs are indexed *Z*_1_, *Z*_2_,…, *Z*_*m*_ in ascending order of *Q*_min_.

To evaluate the efficiency of identity swap discovery, we defined a family of correction effort metrics, denoted 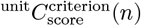. Here, “unit” specifies the basis of measurement (frames or ZODs), “criterion” the stopping rule (e.g., all swaps, threshold-based), and “score” the ranking statistic (here *Q*_min_). First, we consider two complementary unit definitions of a *Q*_min_-based correction effort metric *C*_*Q*_:

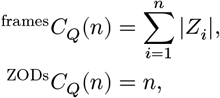

where |*Z*_*i*_| is the number of frames spanned by ZOD *Z*_*i*_. Thus, ^frames^*C*_*Q*_(*n*) represents the cumulative number of frames reviewed after examining the first *n* ZODs, while ^ZODs^*C*_*Q*_(*n*) simply counts the number of ZODs reviewed.

To formalize the notion of the total correction effort required to find all swaps, let 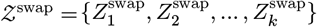 denote the subset of ZODs that contain identity swap instances. A swap-containing ZOD 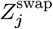 is considered detected if it overlaps with at least one of the first *n* ZODs in the ranking, {*Z*_1_,…, *Z*_*n*_}. The minimal number of ZODs that must be reviewed to detect all swaps is therefore

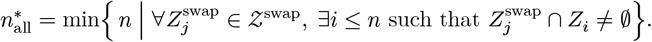

The corresponding correction efforts are

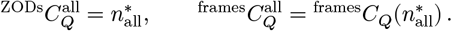

As a complementary criterion, we also quantified the correction effort required to review all ZODs with severity above a fixed threshold. Specifically, let

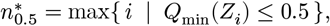

that is, the number of top-ranked ZODs whose minimum quality score does not exceed 0.5. The corresponding correction efforts are

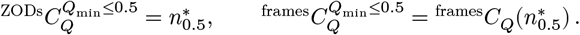

### 6 Alignment of Keypoints to Segmentation Masks

#### SLEAP to Cutie Alignment Algorithm

We aligned SLEAP pose estimation tracks to the stable animal identities defined by consensus Cutie masks on a frame-by-frame basis.

For each frame, we computed the number of SLEAP keypoints that overlapped with Cutie segmentation masks after applying a 5-pixel binary dilation. Dilation was performed on fourfold-downsampled Cutie masks with dimensions 384 × 512 pixels. This improved the robustness of keypoint assignment near animal edges, allowing a boundary keypoint to be associated with multiple adjoining masks rather than risk being incorrectly assigned to only one.

Let *M* = (*M*^(1)^, *M*^(2)^,…, *M*^(*n*)^) denote the set of *n* consensus masks for a given frame, and let *𝒫*_*n*_ denote the set of all *n*! permutations of the index set {1, 2, …, *n*}. As before, for each permutation *π* ∈ *𝒫*_*n*_, we define a permuted version of the masks as

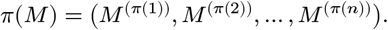

For each keypoint track *x* ∈ {1, …, *n*} at frame *t*, let 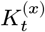 be the number of detected keypoints and let 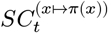 be the number of those keypoints that fall within the dilated mask of Cutie track *π*(*x*). We then defined an alignment score for each permutation *π* as

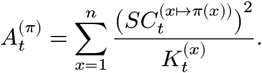

These alignment scores can be interpreted as the product of two intuitive terms. Each fraction 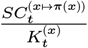 represents the accuracy of assigning SLEAP keypoints from track *x* to Cutie track *π*(*x*), while the multiplier 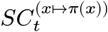 reflects the absolute number of keypoints supporting that assignment. Thus, each summand can be rewritten as

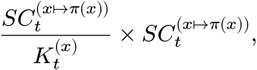

i.e., accuracy × hits. This formulation ensures that both the proportion of correctly aligned key-points (accuracy) and the total number of supporting keypoints (confidence) jointly contribute to the alignment score.

The final alignment at frame *t* was determined by selecting the permutation 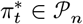 that maximized the alignment score:

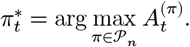

This choice of 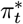 provided the definitive identity mapping between SLEAP tracks and Cutie consensus masks at that frame, thereby resolving the animal identities for the detected keypoint skeletons.

#### Two-Animal Alignment

##### Specialized Definitions

In each male/female interacting dyad, the female vole was assigned to Cutie masks track 0, and the male vole to track 1. SLEAP track numbers were those produced by the SLEAP tracking and track-cleaning pipeline. To evaluate alignment, we defined two permutations *π* ∈ *𝒫*_2_ mapping SLEAP tracks to Cutie tracks:

- **Identity mapping** *π*^ident^: *π*^ident^(0) = 0, *π*^ident^(1) = 1
- **Flipped mapping** *π*^flip^: *π*^flip^(0) = 1, *π*^flip^(1) = 0

The final alignment for each frame was determined by whichever permutation achieved the higher score, 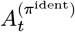 or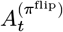. In practice, most frames were definitively aligned by this procedure.

The small minority of non-definitive cases fell into four categories:

- **No keypoints:** frames with no detected SLEAP keypoints.
- **No valid masks:** frames within a double-garbage ZOD, for which there were no valid Cutie segmentation masks from either forward or reverse inference.
- **Zero overlap:** frames with no overlap between SLEAP keypoints and Cutie masks.
- **Ambiguous alignment:** frames in which 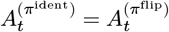, such as when SLEAP keypoints were evenly divided between the two Cutie masks.

##### SLEAP to Cutie Alignment Performance

ZOD resolution and SLEAP–Cutie alignment heuristics enabled us to align 99.4% of SLEAP pose estimation frames with Cutie consensus masks. The remaining 0.58% of frames were primarily concentrated in 149 contiguous video segments across the 83-video dataset, each corresponding to regions where both forward and reverse Cutie mask tracks were rejected during manual review (e.g., double-garbage ZOD). For these mask-free segments, identity swaps were manually corrected using the SLEAP graphical user interface and integrated with the remainder of the dataset, where identity errors were otherwise resolved automatically via Cutie alignment.

SLEAP-to-Cutie alignment statistics.

**SLEAP-to-Cutie alignment statistics**
| Category | Frame Count | Proportion of Total Frames |
| --- | --- | --- |
| Total frames for $n = 83$ videos | 29,880,000 | 1.0 |
| Frames with definitive alignment | 29,703,032 | 0.9941 |
| Frames with no keypoints | 334 | 0.0000112 |
| Frames with no valid masks | 173,129 | 0.00579 |
| Frames with non-overlapping keypoint/mask data | 55 | 0.00000184 |
| Frames with ambiguous alignment | 3,450 | 0.000115 |

Double-garbage ZOD (dgZOD) statistics.

**Double-garbage ZOD (dgZOD) statistics**
| Metric | Value |
| --- | --- |
| dgZOD count | 149 |
| dgZOD total frames | 172,980 |
| Mean dgZOD frames | 1,160.94 |
| Median dgZOD frames | 239 |
| Minimum dgZOD length | 11 |
| Maximum dgZOD length | 12,922 |

##### SLEAP Tracking Error Estimation

To detect potential SLEAP identity swaps, we monitored changes in the relative magnitude of the alignment scores 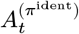 and 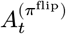. A change in sign—where 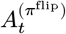 exceeds 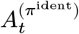 after previously being lower—was flagged as a candidate identity swap. If the immediately preceding frame was successfully aligned using Cutie masks, the swap was localized to the current frame. However, if the preceding frame fell within a gap of frames that could not be aligned (e.g., due to missing or invalid Cutie masks), the precise location of the swap could not be determined and the event was excluded from downstream analyses of SLEAP identity swaps.

## 7 Tools Availability

We make available tools for (1) keyframe generation and proofing using the Segment Anything Model, (2) video object segmentation using our bidirectional scheme, (3) 2-animal ZOD detection, annotation, and review, and (4) alignment of keypoints to segmentation masks. The code repository will be hosted at github.com/bondscape/bidirectionalVOS.

## 8 Computing Resources

We performed Cutie VOS on both desktop and HPC cluster environments running Ubuntu 20.04 and Rocky Linux 8.10, respectively. The minimal tested desktop configuration included 32 GB RAM and an NVIDIA RTX 2080 Ti GPU with 11 GB VRAM. On the HPC cluster, jobs were submitted with a minimum GPU memory request of 10 GB. Performance on lower-specification machines was not evaluated.

The software environment included Python 3.11.5, `opencv-python` 4.8.1.78, `pytorch-cuda 12.1`, and `pytorch py3.11_cuda12.1_cudnn8.9.2_0`.

Our bidirectional VOS pipeline comprised three key phases: (1) FFmpeg-based generation of video clips from a full-length recording, (2) FFmpeg-based reversal of clips, and (3) Cutie VOS inference. The duration of clip generation scaled with the frame count of the full-length video, and reversal required 1.5 minutes for 15,000 frames. Cutie inference averaged 12–14 frames per second for videos at 2048 × 1536 px resolution.

## Acknowledgements

We would like to thank members of the Manoli lab and Kevin Bender for their thoughtful feedback on figures.

## Author contributions

S.W. and C.K. conceptualized the project. S.W. and K.Q. developed the methods. K.Q. produced the codebase. A.J. and S.W. collected behavior videos. S.W. and S.D. curated behavior data. S.W. performed the analyses. S.W. wrote the manuscript, with feedback from K.Q., D.S.M., and C.K.

## Data and materials availability

Codebase will be made available on Github.

## Supplementary Materials

Supplementary Figure 1: SLEAP identity swaps occur throughout long recordings and are difficult to identify using post hoc keypoint-based metrics.

Supplementary Figure 2: User-guided tools for generating keyframe masks and resolving segmentation mismatches.

Supplementary Figure 3: Comparison of manual review effort and identity swap resolution between SLEAP and Cutie across an 83-video dataset.

Supplementary Figure 4: Schematic of segmentation swap types and their corresponding IOU values.

Supplementary Figure 5: Distribution of mask permutation IOU values for three-animal tracking.

