## Supplementary Figures for "Identity-stable multi-animal tracking using bidirectional segmentation with object-level memory"

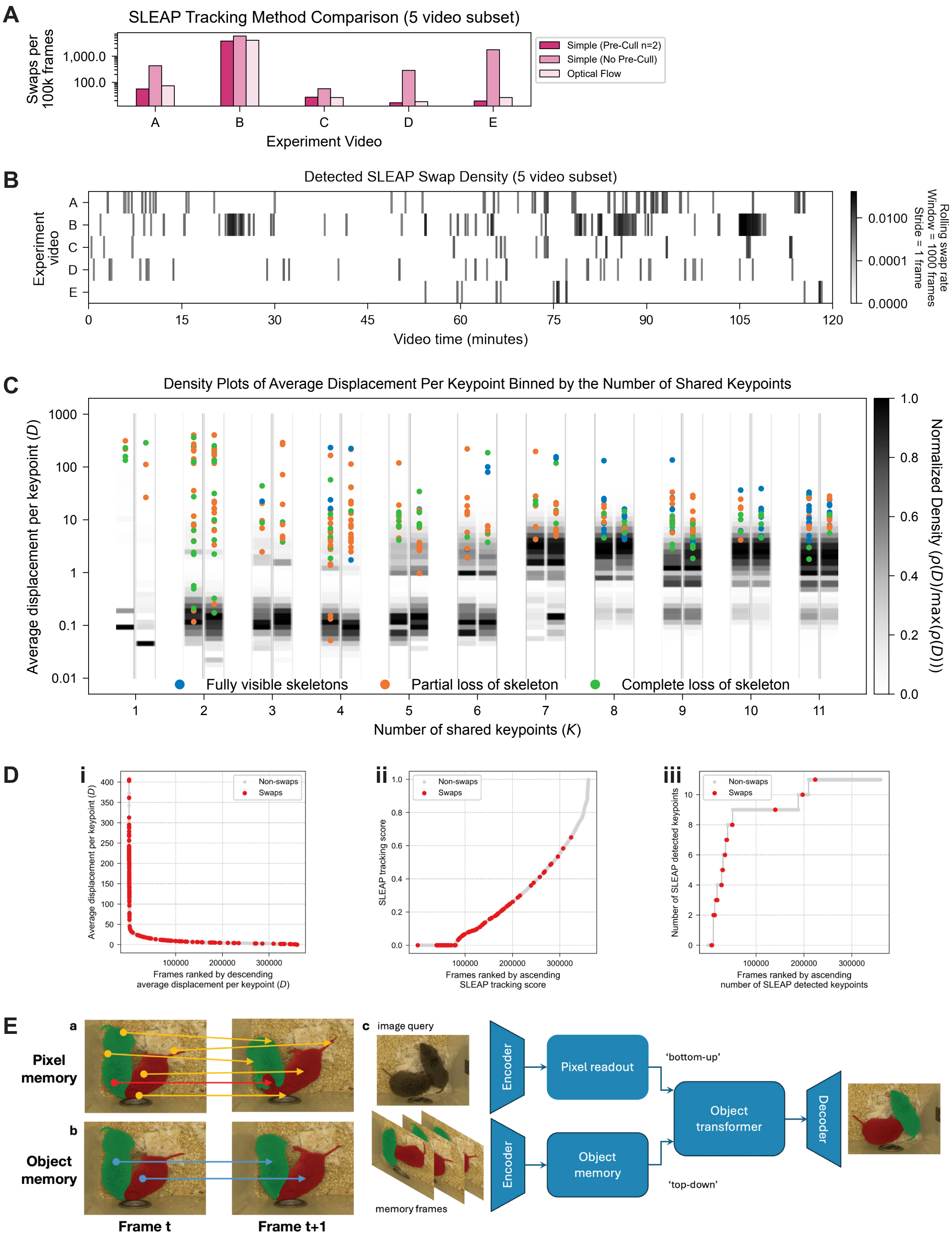


**Supp. Figure 1: SLEAP identity swaps occur throughout long recordings and are difficult to identify using post hoc keypoint-based metrics.**

A. Performance comparison of three post hoc tracking strategies across a subset of five manually annotated two-hour videos. First, swaps were manually identified from the *simple (pre-cull to two instances)* tracking method. Using those annotations as ground truth, we then computationally annotated swaps for the *simple (no pre-cull)* and *optical flow* tracking methods. The *simple (no pre-cull)* method produced substantially higher swap rates, while the *simple (pre-call to two instances)* and *optical flow* methods yielded similar results. All subsequent figures use keypoint data from the *simple (pre-cull to two instances)* method for efficiency.

B. Swap density across the five manually annotated videos, plotted using a 15-second rolling average. In most cases, identity swaps occur throughout the recordings.

C. Relationship between motion energy, the number of keypoints detected, and swap occurrence in the single representative video shown in Fig. 1C. Motion energy is defined as the normalized Euclidean distance between shared keypoints in adjacent frames. Although swaps are enriched at high motion energy, many fall within the main density peak, limiting its discriminative value. Additionally, when few keypoints are detected, low motion energy can also be associated with swaps. (Color coding matches Fig. 1A and 1B.)

D. Ranked metric curves for the representative video shown in Fig. 1C. For each pair of adjacent frames, we generated three candidate metrics for identity swap detection: (i) motion energy, (ii) SLEAP tracking score, and (iii) number of detected keypoints. Frame pairs were then rank-ordered by each metric, and swap positions were overlaid. Although swaps are enriched toward the top of each ranking, there are swaps scattered throughout the tails of the curves, underscoring the limited reliability of these metrics for producing identity error-free data in an efficient manner. See Fig. 3B for AUROC statistics across the full dataset of 83 videos.

E. Schematic of Cutie, a query-based video object segmentation algorithm developed by Cheng *et al*.[47] Cutie leverages both bottom-up pixel memory and top-down object-level memory to promote identity continuity over time.

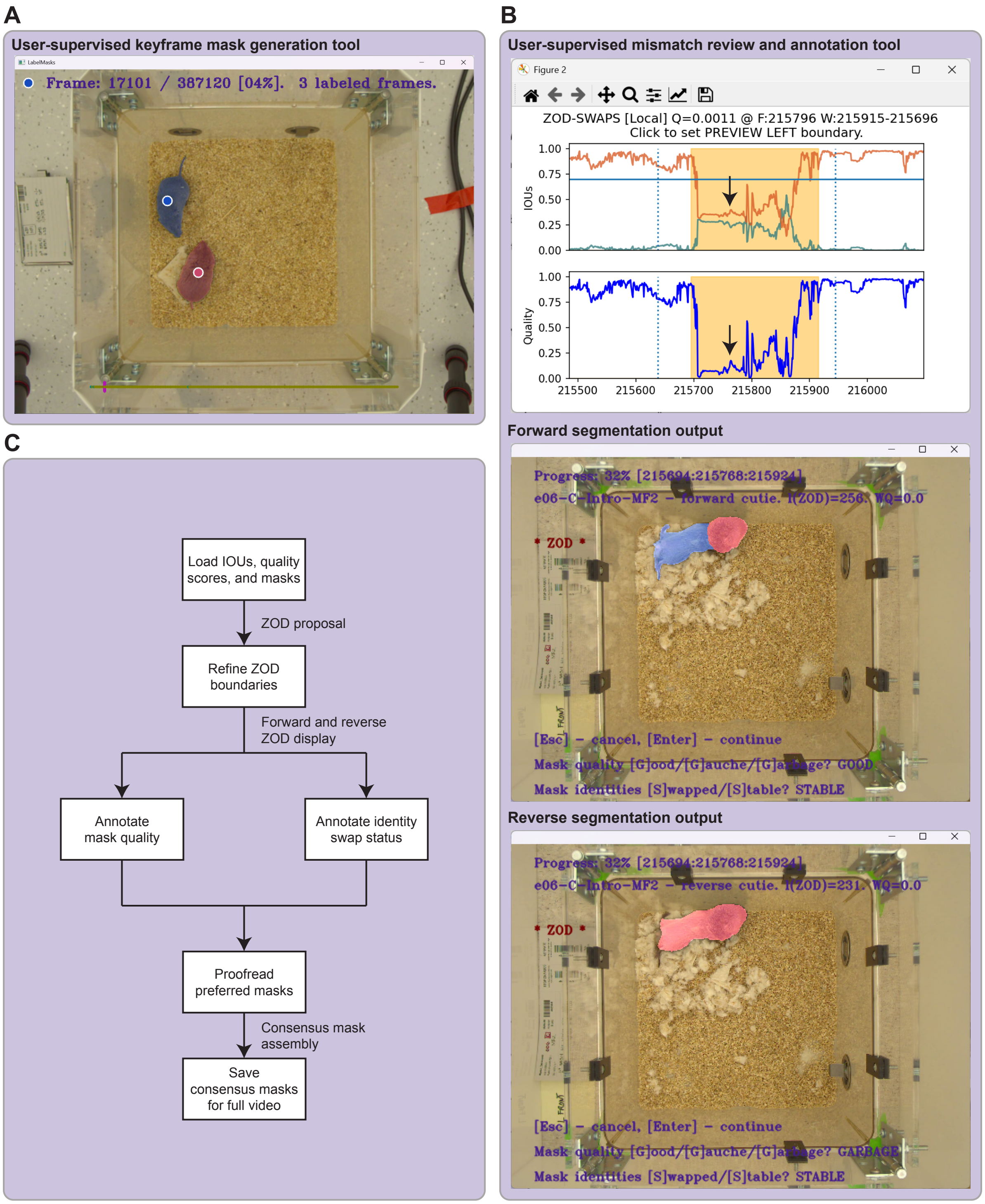


**Supp. Figure 2: User-guided tools for generating keyframe masks and resolving segmentation mismatches.**

A. Keyframe mask generation process. Users designate the two voles with white-rimmed blue and red dots (order arbitrary). Optional negative prompts can be added to suppress background objects (not shown). Segment Anything uses the user-defined prompts to generate candidate masks, and users select the best of three outputs for each animal.

B. ZOD (zone of disagreement) annotation interface. Top: Interactive GUI displaying framewise IOU $\pi^{ident}$ (teal), IOU $\pi^{flip}$ (burnt orange), and quality scores (blue). Users define ZOD boundaries to span regions flanked by high-quality masks. A butterscotch-shaded region highlights the selected ZOD. The black arrow indicates the position of the frame whose forward and reverse segmentation outputs are shown below. For each ZOD, users independently review forward (middle) and reverse (bottom) segmentation outputs and assess (1) whether identity is preserved across the segment and (2) whether the masks are high-quality, suboptimal but usable, or unusable altogether. In this example, reverse inference erroneously produced a single merged mask spanning both animals (with the second mask missing), whereas forward inference yielded two accurate masks. Forward segmentation is therefore retained for consensus.

C. Flowchart illustrating user-driven (in boxes) and automated (beside arrows) steps in the annotation process for bidirectional VOS.

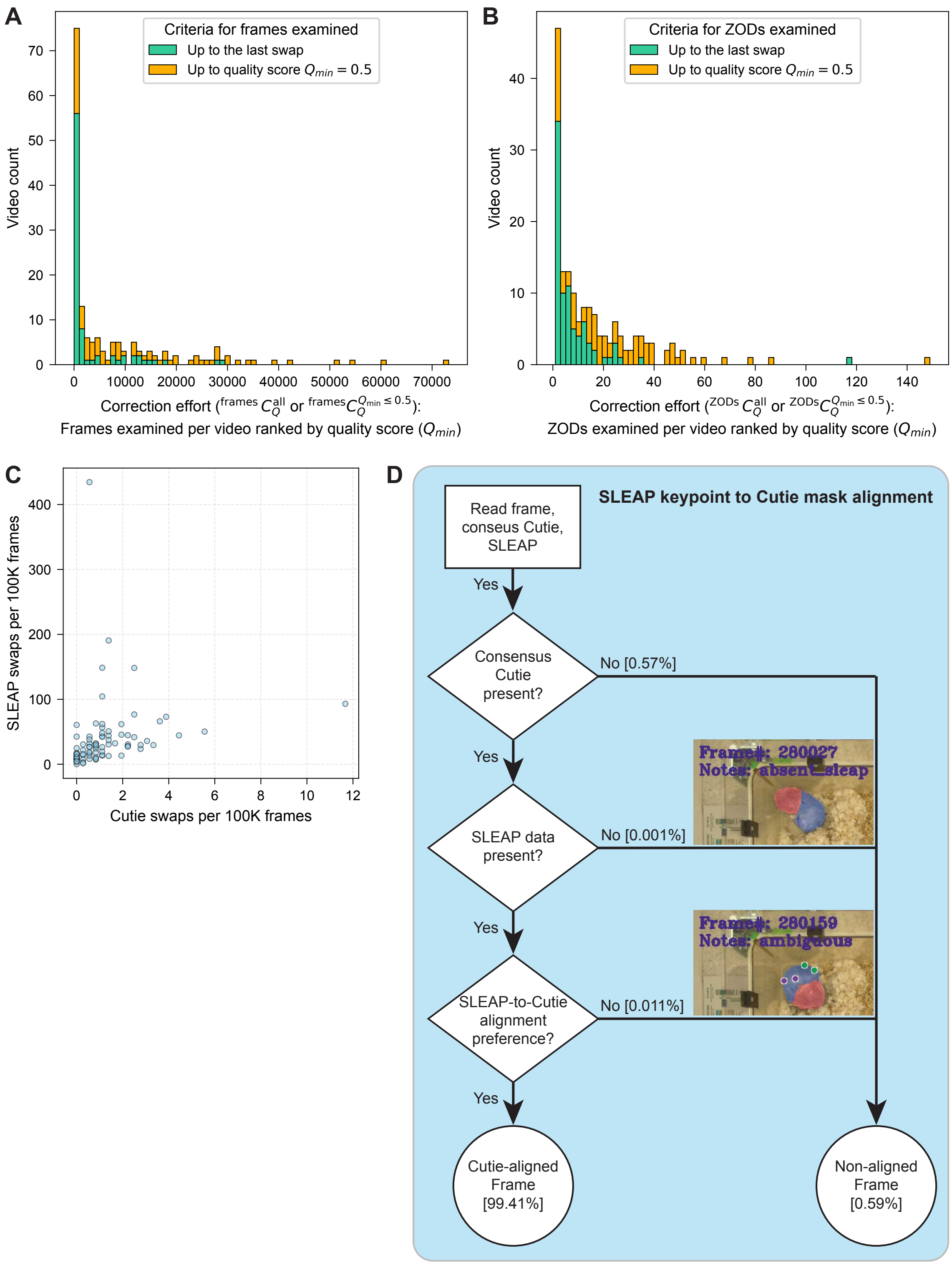


**Supp. Figure 3: Comparison of manual review effort and identity swap resolution between SLEAP and Cutie across an 83-video dataset.**

A. Number of frames manually reviewed to recover all Cutie identity swaps (mint green) versus the number needed to reach a quality score threshold of 0.5 (golden), which was sufficient to detect swaps across all videos. Bars show the distribution across 83 videos.

B. Number of ZODs reviewed to uncover all Cutie identity swaps (mint green) versus the number needed to reach the quality score threshold of 0.5 (golden).
C. Scatterplot comparing the total number of identity swaps detected using SLEAP versus Cutie for each video. Note the differing axis scales.
D. Flowchart illustrating the alignment procedure for assigning SLEAP track identities to Cutie-derived consensus identities. Percentages indicate the fraction of frames across all 83 videos (360,000 frames each) falling into each outcome category. Two rare scenarios – absence of keypoint detections and ambiguous identity assignment – are shown as examples.

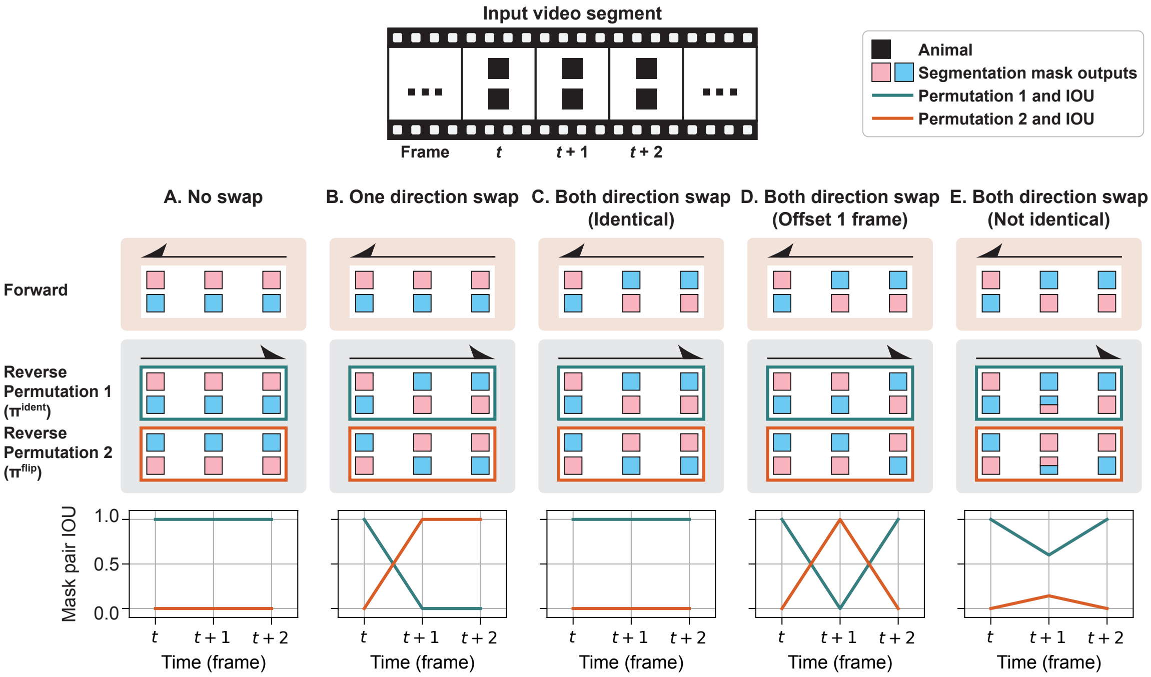


**Supp. Figure 4: Schematic of segmentation swap types and their corresponding IOU values.**
Forward and reverse segmentation masks are simulated for three consecutive frames from an input video segment. The teal box and curve reflect IOU comparisons between forward masks and reverse masks permutation 1, while the burnt orange box and curve reflect equivalent values for permutation 2.

A. No-swap baseline.
B. A swap in one direction but not the other produces a characteristic X-shaped crossing of IOU curves (see Fig. 2C).
C. Swaps in both directions with identical pixel-level error show no visible IOU curve deviations; in practice, this has not been observed due to the history-dependent trajectories of forward and reverse segmentations.
D. Swaps in both directions that are offset by a single frame produce a sharp crossing of IOU curves, flagging the interval as a ZOD.
E. Coinciding but only partially overlapping bidirectional swaps can cause more subtle IOU deflections, requiring careful selection of the IOU threshold for examination.

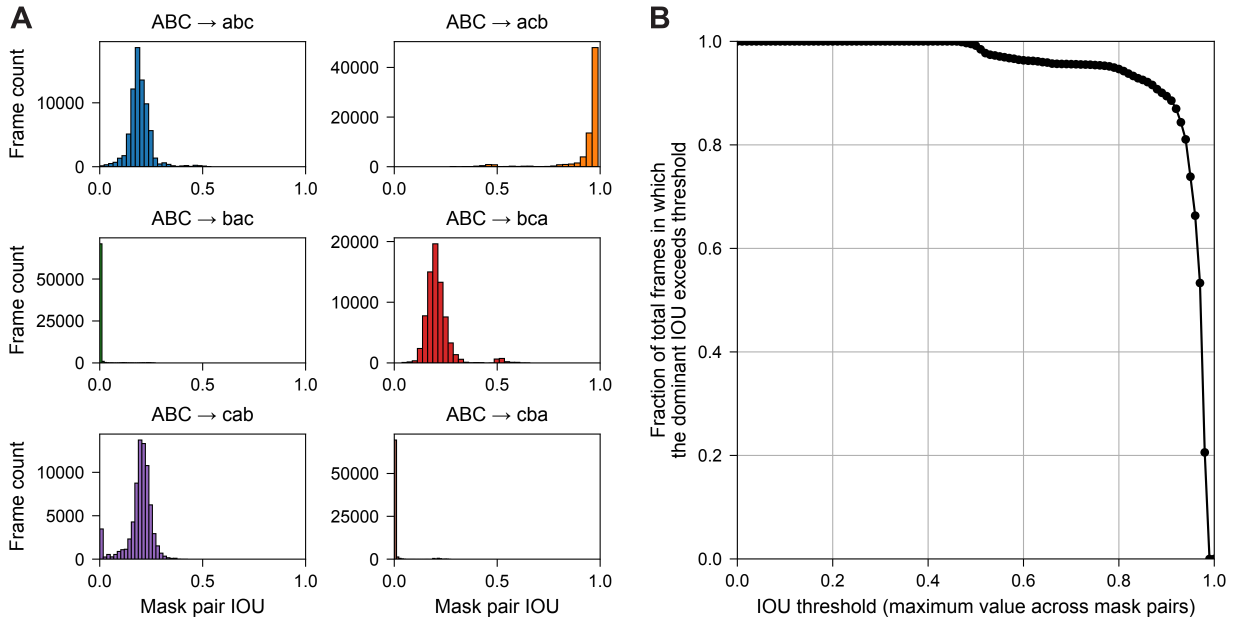


**Supp. Figure 5: Distribution of mask permutation IOU values for three-animal tracking.**

A. Histograms of IOU values for each of the six possible forward-to-reverse mask mappings, corresponding to the IOU curves in Fig. 4B.
B. Fraction of total frames (y-axis) in which the dominant IOU – the highest of the six permutation IOUs for a given frame – exceeds a given IOU threshold (x-axis).
